# RNA-sequencing indicates immune cell signaling and inflammatory gene expression in cardiac fibroblasts increases with developmental age

**DOI:** 10.1101/2021.03.01.433442

**Authors:** Luke R. Perreault, Thanh T. Le, Madeleine J. Oudin, Lauren D. Black

## Abstract

**Background:** Cardiac fibroblasts are responsible for extracellular matrix turnover and repair in the cardiac environment and serve to help facilitate immune responses. However, it is well established that they have significant phenotypic heterogeneity with respect to location, physiological conditions, and developmental age. The goal of this study was to provide an in-depth transcriptomic profile of cardiac fibroblasts derived from rat hearts at fetal, neonatal, and adult developmental ages to ascertain variations in gene expression that may drive functional differences in these cells at these specific stages of development.

**Results:** **We performed** RNA-seq of cardiac fibroblasts isolated from fetal, neonatal, and adult rats was performed and compared to the rat genome. Principal component analysis of RNA-seq data suggested data variance was predominantly due to developmental age. Differential expression and Gene set enrichment analysis against Gene Ontology and Kyoto Encyclopedia of Genes and Genomes datasets indicated an array of differences across developmental ages, including significant decreases in cardiac development and cardiac function-associated genes with age, and a significant increase in immune and inflammatory-associated functions - particularly immune cell signaling, and cytokine and chemokine production - with respect to increasing developmental age.

**Conclusion:** These results reinforce established evidence of diverse phenotypic heterogeneity of fibroblasts with respect to developmental age. Further, based on our analysis of gene expression, age-specific alterations in cardiac fibroblasts may play a crucial role in observed differences in cardiac inflammation and immune response observed across developmental ages.

## Introduction

While cardiomyocytes (CMs) have historically been the therapeutic target for bioengineering approaches to heart repair (1–3), non-cardiomyocyte cell populations in the heart often play a critical role in the heart’s response to injury and, as such, have been subject to increased appreciation and study over the past few decades. Non-myocyte cell populations, which include vascular cells, endothelial cells, immune cells, and cardiac fibroblasts (CFs), comprise approximately 70% of the total cell population in the adult mammalian heart (4–6), and CFs are the major fraction of these non-myocyte cells. CFs, in particular, play a major role in mediating cardiac homeostasis, facilitating the development and maturation of cardiac ECM and promoting cardiomyocyte proliferation *in utero* (7), and yet they are also integral in the initiation and progression of various cardiac pathologies (8–11).

CFs are classically defined as mesenchymal-like cells responsible for extracellular matrix (ECM) production and remodeling (11). These cells are largely quiescent in the adult mammalian heart but exhibit extensive phenotypic plasticity, able to react to stimuli following injury and transition to a myofibroblast phenotype. In contrast to CFs in a resting state, myofibroblasts are highly active, migrating into and proliferating rapidly in the wound environment, where they extensively upregulate collagen I production to produce mature fibrotic scar tissue in adult mammals (12, 13). However, the role of CFs in responding to injury extends beyond ECM regulation and fibrosis. CFs are an important immune mediator and contributor to the inflammatory process (14, 15). One study by Kawaguchi *et al* indicated that mouse CFs exposed to lipopolysaccharide (LPS), an inflammatory stimulus, promoted interleukin-1β (IL-1β, a major inflammatory mediator) expression in those cells, but not in CMs. The CFs were found to act as “sentinel” cells in the heart, deploying a broad range of chemokines including MCP-1, macrophage inflammatory protein-1, and RANTES (15) to aid immune cell influx into a wound environment (16).

Despite their critical roles in the heart, CFs are loosely defined with respect to their phenotype, which varies considerably across developmental ages. The cell type traditionally considered a fibroblast is now understood to be a heterogeneous population of cells arising from different progenitors (17, 18), with a lack of specific markers to establish their identity *in vitro,* although CFs from both fetal and adult sources express periostin, vimentin, and discoidin domain receptor 2 (DDR2) (19). Further, through crosstalk with cardiomyocytes, CFs influence cardiac response to injury differently with respect to developmental age. Fetal CFs are implicated in driving cardiomyocyte growth and proliferation via secretion of paracrine factors, whereas adult CF-secreted factors are implicated in inducing cardiac hypertrophy, findings that reflect the regenerative healing observed in fetal and neonatal hearts, and fibrosis and hypertrophy observed in injured adult cardiac tissue (20, 21). As there is a significant, rapid shift in ECM composition (22, 23) and mechanical function of the heart pre- to post-birth, studying the phenotype of neonatal CFs would also be valuable, although there is scarce literature currently available on CFs of this specific developmental age. A single-cell RNA-seq study has established that transcriptomic changes in mouse CFs from a neonatal to adult state promote concomitant CM maturation, connecting age-specific CF phenotypic changes with cardiac maturation (24).

Studies of CF behavior across various developmental ages are invaluable to further elucidating the identities and functions of this heterogeneous cell population and how it changes with age, particularly with respect to the healing response of cardiac tissue. Many analyses have investigated how CF developmental age impacts cardiac structure and function or induces functional change in CMs (3, 25). By contrast, an understanding of how CF gene expression related to immune infiltration and the inflammatory response change with age is not well-established, despite the critical roles of these cells in cardiac inflammation and immunity and how critical these processes are to the healing response (15, 26). In this study, we employ RNA-seq transcriptomic analysis to evaluate rat cardiac fibroblast gene expression at fetal, neonatal, and adult developmental ages. This transcriptomic profile presents a broad view of CFs across development and aging to facilitate age-specific analyses of these cells. However, we specifically investigate age-dependent changes in immune and inflammatory gene expression, speculating that CF involvement in these processes increases with development. We found that compared to postnatal day 1 neonatal CFs, fetal CFs display significantly downregulated immune- and inflammatory-implicated genes and pathways, whereas adult CFs markedly increased expression of immune and inflammatory proteins. Taken together, this suggests CFs adopt an inflammatory/immune-support role rapidly after birth, that is further enhanced with continued development.

## Results

### Fibroblast sequencing, culture, and characterization

Cardiac fibroblasts (CFs) were isolated from fetal (E18), neonatal (P1), and young adult (~7-week-old, at which point the rats are sexually mature (27)) rat hearts. After pre-plating the isolated mixed cardiac cells (28, 29), isolates were cultured to confluency and used for RNA-sequencing or expanded out to passages 2-4 for additional experiments and culture validation (Fig. 1).

**Figure 1:**
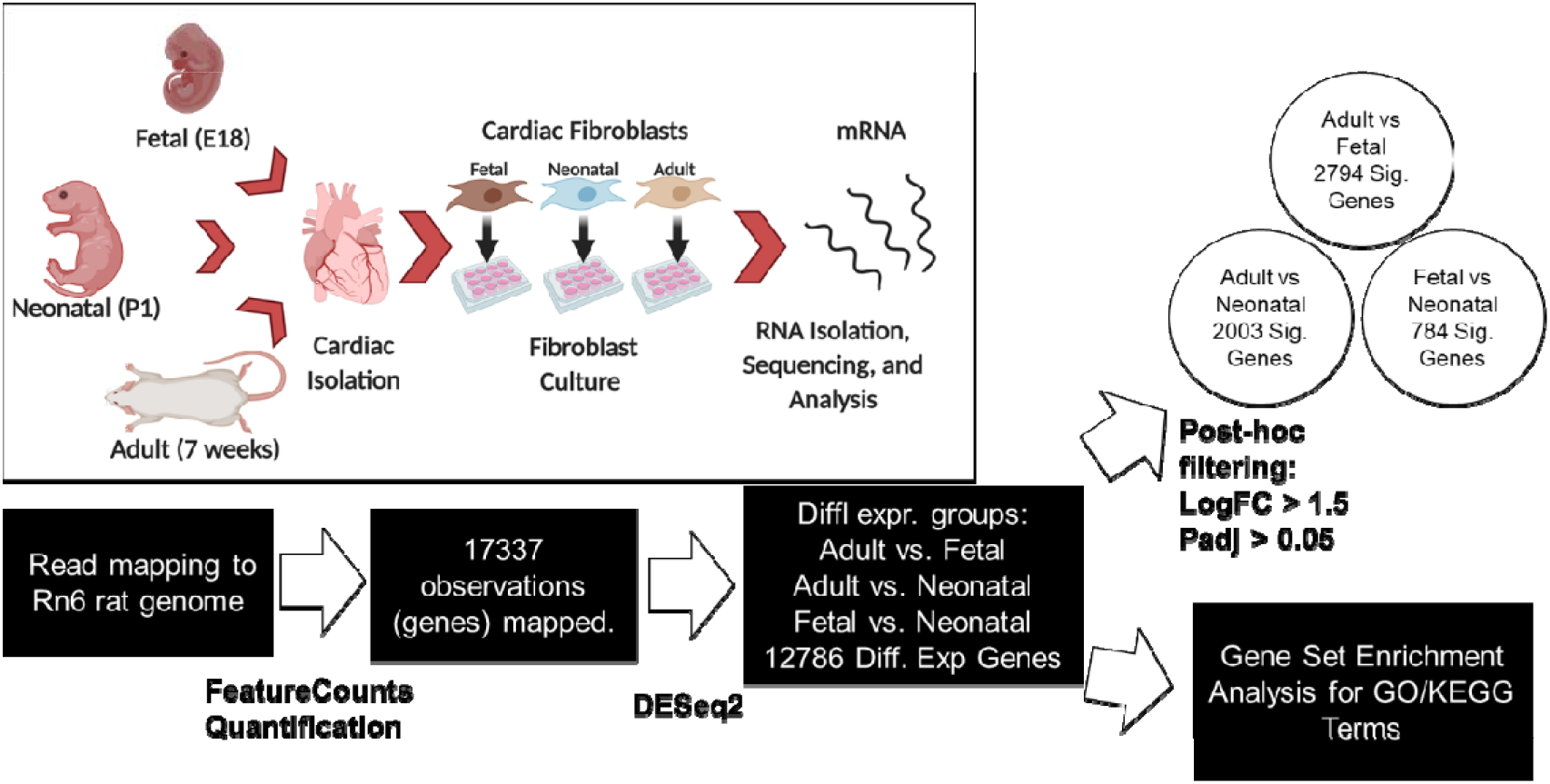
Pipeline for RNA isolation and differential expression analysis. RNA is mapped to Rn6 rat genome via STAR and reads counted, yielding 17337 distinct observations. These are normalized and used in differential expression analysis in DESeq2, and then used directly in GSEA for enrichment analysis, or filtered via LogFC and adjusted p-value to evaluate significantly expressed genes. Insert: Schematic depicting isolation of three distinct ages of rat cardiac fibroblasts, followed by culture and RNA isolation. Schematic creating using art acquired from BioRender.

RNA-seq mapping to the Rn6 rat genome yielded a total of 17337 discrete genes mapped. Differential expression analysis yielded a total of 12786 genes of interested between comparison conditions (adult vs fetal, adult vs neonatal, and fetal vs neonatal) (Fig. 1).

CFs grown in culture were imaged following immunofluorescence labeling and fibroblast lineage was confirmed by demonstration that the cells were negative for markers of endothelial (CD31) (Fig 2D-F) or hematopoietic (CD45) lineages (Fig 2G-I), respectively (30, 31). Cells were positive for vimentin (Fig 2A-C), a broad CF marker (31, 32). Cultured fibroblasts all displayed similar elongated morphology and positive vimentin expression throughout. (Fig 2).

**Figure 2:**
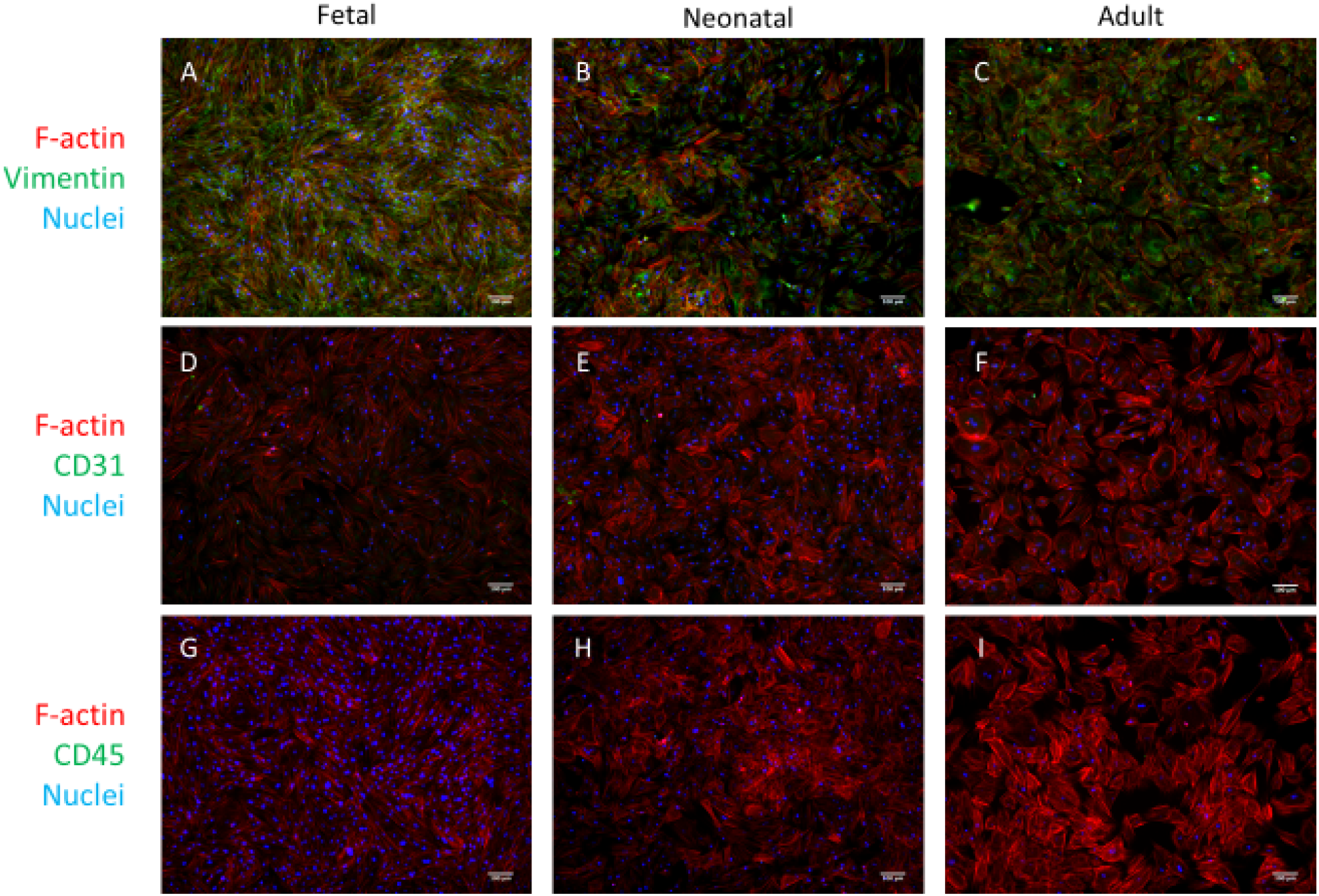
Immunofluorescence microscopy characterization of primary P2-P4 cardiac fibroblast cultures. A-C) Vimentin staining of fetal (A), neonatal (B) and adult (C) cells, in order to identify cardiac fibroblasts. D-F) CD-31 staining of fetal (D), neonatal (E), and adult (F) cells to identify endothelial-lineage cells in culture. G-I) CD-45 staining of fetal (G), neonatal (H), and adult (I) cells to identify hematopoetic-lineage cells in culture. Scale bars = 100um.

### Clustering analysis reveals broad variances in gene expression based on CF age

Sequencing data of fetal, neonatal, and adult CFs (groups referred henceforth as FCF, NCF, and ACF, respectively) were analyzed and normalized before differential expression analysis was executed, all in the *DESeq2* package in R. Sample-to-sample Manhattan distance was computed and a heatmap showing sample distance and correlation was produced (Fig 3A).

**Figure 3:**
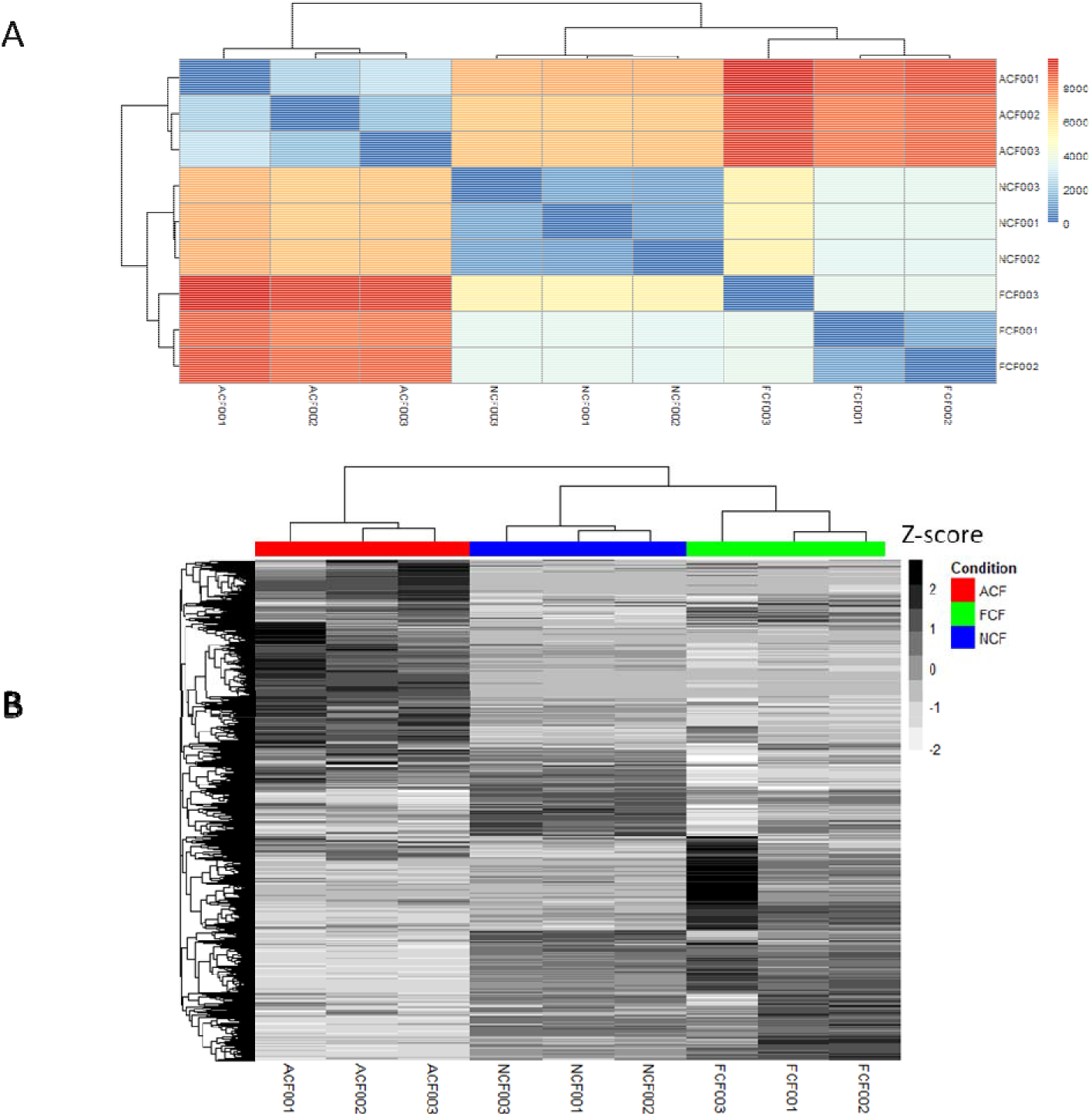
Normalized RNA-seq sample data from adult cardiac fibroblasts (ACF), neonatal cardiac fibroblasts (NCF) and fetal cardiac fibroblasts (FCF). A) Sample-to-sample distance heatmap utilizing Manhattan-distance calculation, showing close clustering of FCF and NCF samples. Heatmap produced in pcaExplorer in R. B) Heatmap of normalized gene counts from all identified genes, showing clustering of genes the three age groups. Genes are row-normalized, and heat-map color is based on z-score. Heatmap produced with R software.

The heatmap indicated a stronger correlation between the FCF and NCF samples. FCF samples had the lowest level of similarity to ACF samples, followed by NCFs, suggesting the sample variability was strongly age dependent. This was supported by unsupervised hierarchical clustering of the DESeq2-normalized geneset across all samples, visually presented in the heatmap in Fig. 3B. FCFs and NCFs clustered together, whereas ACFs diverged greatly in their gene expression compared to the two younger CF sources.

Principal component analysis (PCA) was evaluated with respect to samples (Fig 4A) and genes (Fig 4B-D). The sample PCA supported the hypothesis that variation in samples was primarily due to age, as it indicated sample distribution along the first principal component (constituting approximately 92% of total variance in the data set) coincided with the associated with age of the fibroblast sample source (Fig. 4A).

**Figure 4:**
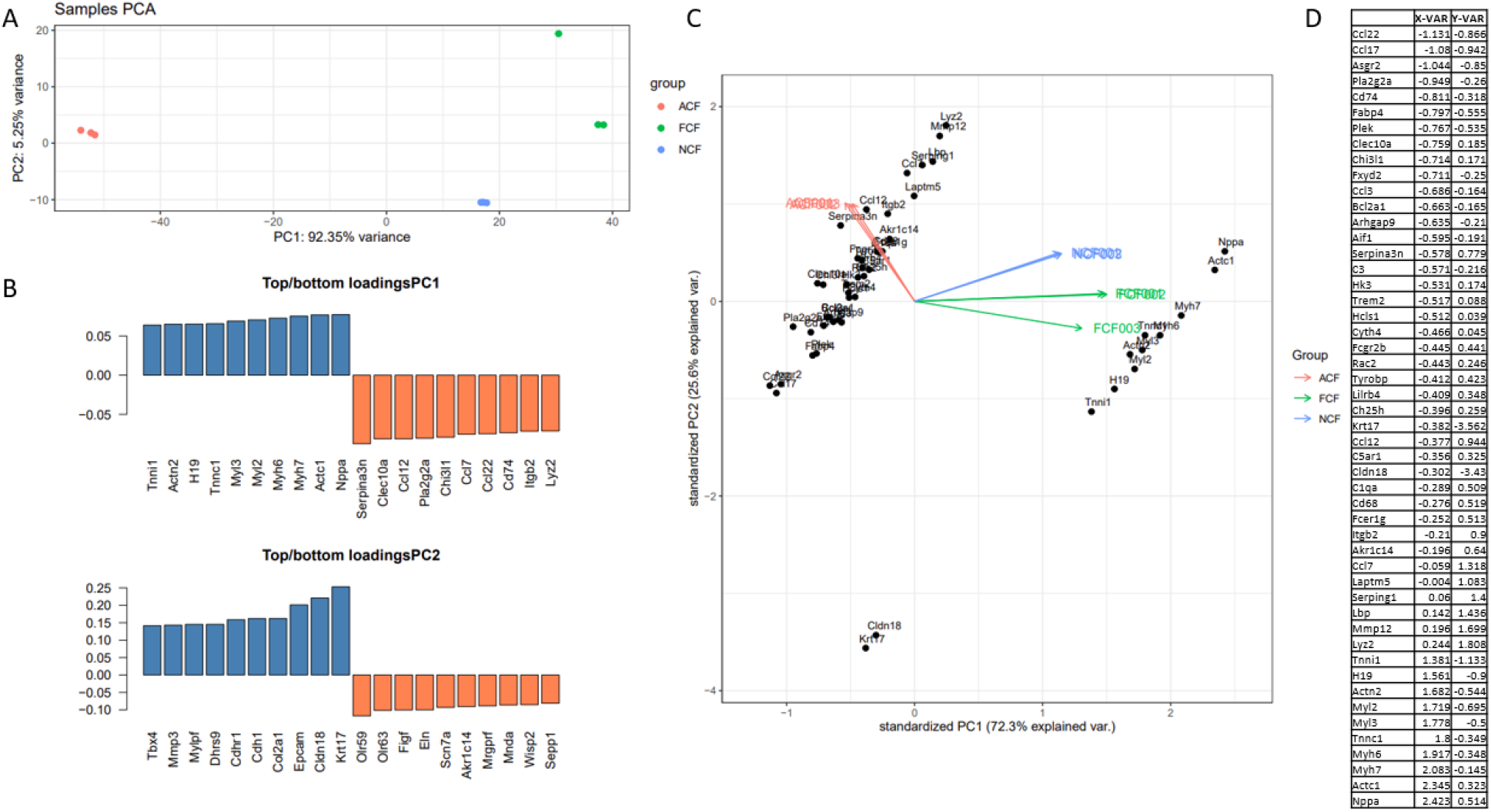
Principal component analysis with associated loadings with respect to gene expression in fetal, neonatal, and adult rats. A) Table of 50 top contributing genes to the PCA analysis, with associated x-coordinates and y-coordinates on the associated PCA graph in (B). B) PCA of top 50 genes contributing most variance to the first two principal components, showing relationship and location with respect to fetal (FCF), neonatal (NCF), adult (ACF) samples. C) Loadings plot for top/bottom ten highest contributing genes of first and second principal components. D) PCA of samples with respect to first two principal components, showing broad distribution with respect to age on the first PC axis. All graphs were generated in pcaExplorer package in R.

Analysis of the 50 genes that most significantly contributed to the PCA variance produced two distinct clusters with respect to the first principal component, which appeared to be associated with CF age. This suggested greater variance between ACFs and the two juvenile CF groups (FCF and NCF), with the juvenile groups associated with genes that clustered closer together. Genes associated with the ACF cluster were broadly immune associated, including chemokines *Ccl22 Ccl17,* and *Ccl3,* and cell surface receptor *Cd74.* The NCF/FCF cluster was observed to be predominantly associated with genes related to cardiac and embryonic development, including *Myh6, Myh7, Nppa, Myl2, Myl3,* and *Actc1* (Fig 4B-D).

Differential expression analysis further supported the transcriptomic variance between adult CFs and neonatal and fetal CFs: post-hoc filtering (log_2_FC > 1.5, padj > 0.05) produced 2794 significant differentially expressed genes in ACFs vs FCFs and 2003 significant differentially expressed genes in ACFs vs NCFs, versus 784 genes differentially expressed between FCFs and NCFs (Fig. 1). The top 25 most significantly upregulated and downregulated genes from each comparison are presented in Table 1.

**Table 1:**
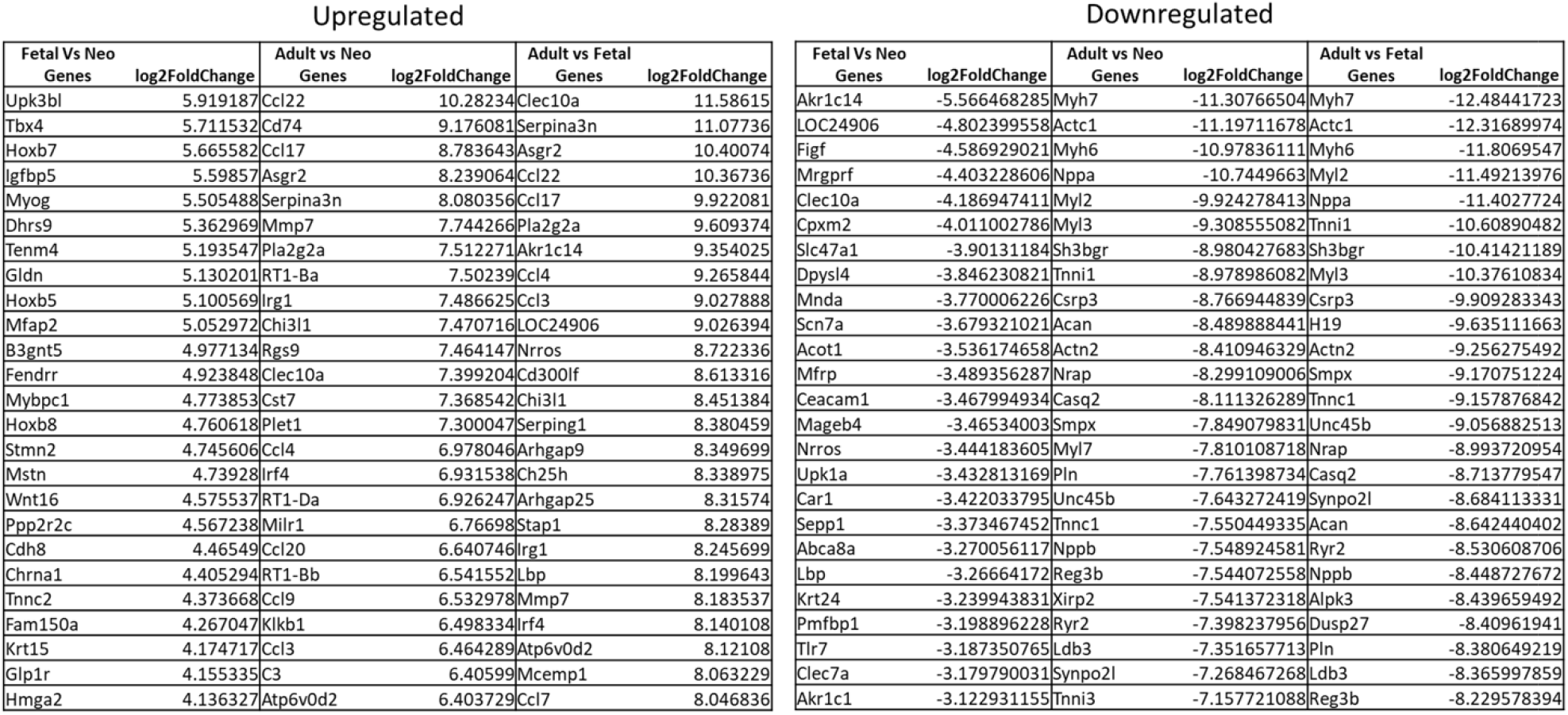
List of 25 most significantly upregulated and 25 most significantly downregulated genes from Fetal vs. Neonatal, Adult vs. Neonatal, Adult vs. Fetal comparisons.

Additional descriptions of functions of the top 10 most significantly upregulated and downregulated genes from each comparison are presented in Additional File 1. A preliminary investigation of gene distribution within these comparisons was conducted using volcano plots generated in the *EnhancedVolcano* R package, using a logFC cutoff of 1.5 and p-value cutoff of 0.001 as parameters. The plot of FCFs vs NCFs indicated that immune-associated genes such as *Ifitm1* and *Lbp,* and matrix regulation-associated genes such as *Ctsc, Figf,* and *Itgb3* were downregulated in FCFs compared to NCFs, whereas developmental genes such as *TBX4, Mdk, Hoxb7,* and *Hoxb8* were upregulated (Fig. 5A, Table 1). ACF comparisons to NCFs and FCFs yielded similar gene distributions in the volcano plots (Fig. 5B-C). In both comparisons, cardiac development-associated genes such as *Actc1, Actg2, Tnnc1,* and *Myh7* were downregulated in ACFs versus the younger group, while immune- and inflammatory-associated genes such as *Tgfb1, Ccl3,* and *Ccl4* were observed in ACFs to be comparatively upregulated (Fig. 5B & 5C, Table 1).

**Figure 5:**
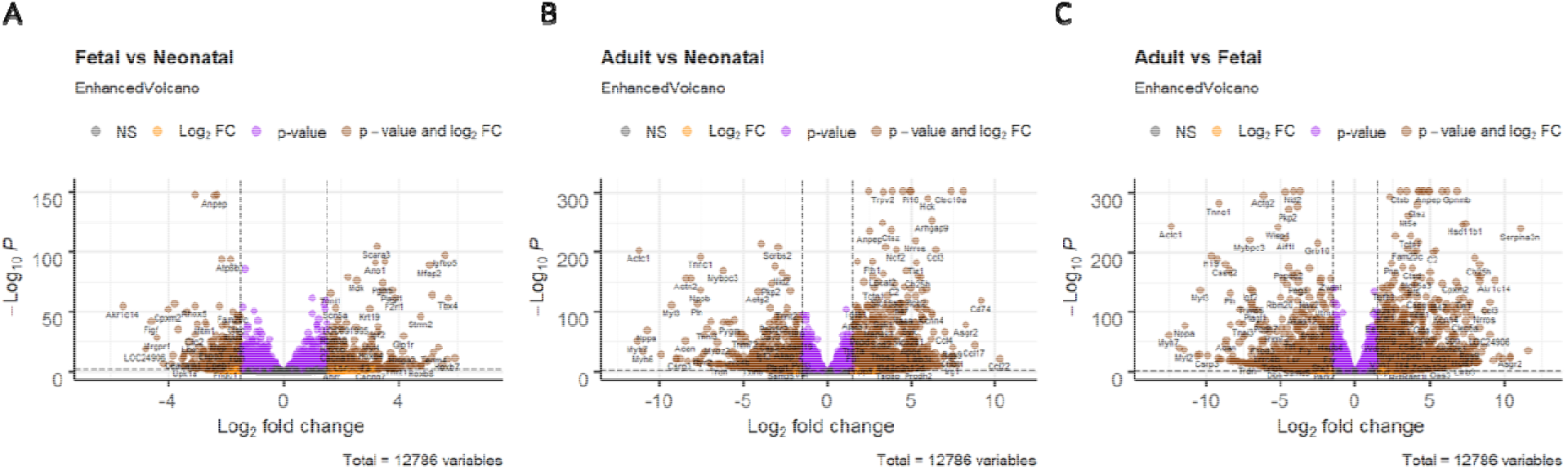
Volcano plots of significant differentially expressed genes. A) fetal vs neonatal, B) adult vs neonatal, C) adult vs fetal, with boundaries set at logFC of 1.5, and p-value of 0.001. Genes color-coded yellow indicate significant LogFC but not statistically significant. Purple genes indicate statistical significance but genes under absolute value of LogFC threshold. Brown color-coded genes meet LogFC threshold and statistical significance. Volcano plots generated in EnhancedVolcano package in R.

### Differential expression indicates age-specific shifts in immune signaling and cardiac development

Differentially expressed gene lists from the three separate comparisons (FCF vs NCF, ACF vs NCF, ACF vs FCF) were ranked based on fold change, and subsequently analyzed using Gene Set Enrichment Analysis (GSEA) methodology (33), within the *clusterProfiler* package in R (34). The full lists of differentially expressed genes identified in these analyses are available in Additional File 3 (Fetal vs Neonatal), Additional File 5 (Adult vs Neonatal), and Additional File 7 (Adult vs Fetal). These datasets were subjected to analysis utilizing Gene Ontology (GO, analyzing for Biological Process, Cellular Component, and Molecular Function terms) (35) and Kyoto Encyclopedia of Genes and Genomes (KEGG) curated datasets (36). Analysis of fetal CFs vs neonatal CFs against GO datasets yielded an array of significantly downregulated ontologies associated with immune response and immune cell signaling (Fig 6A, Table 2), including “leukocyte mediated immunity,” “positive regulation of immune response,” and “activation of immune response.”

**Figure 6:**
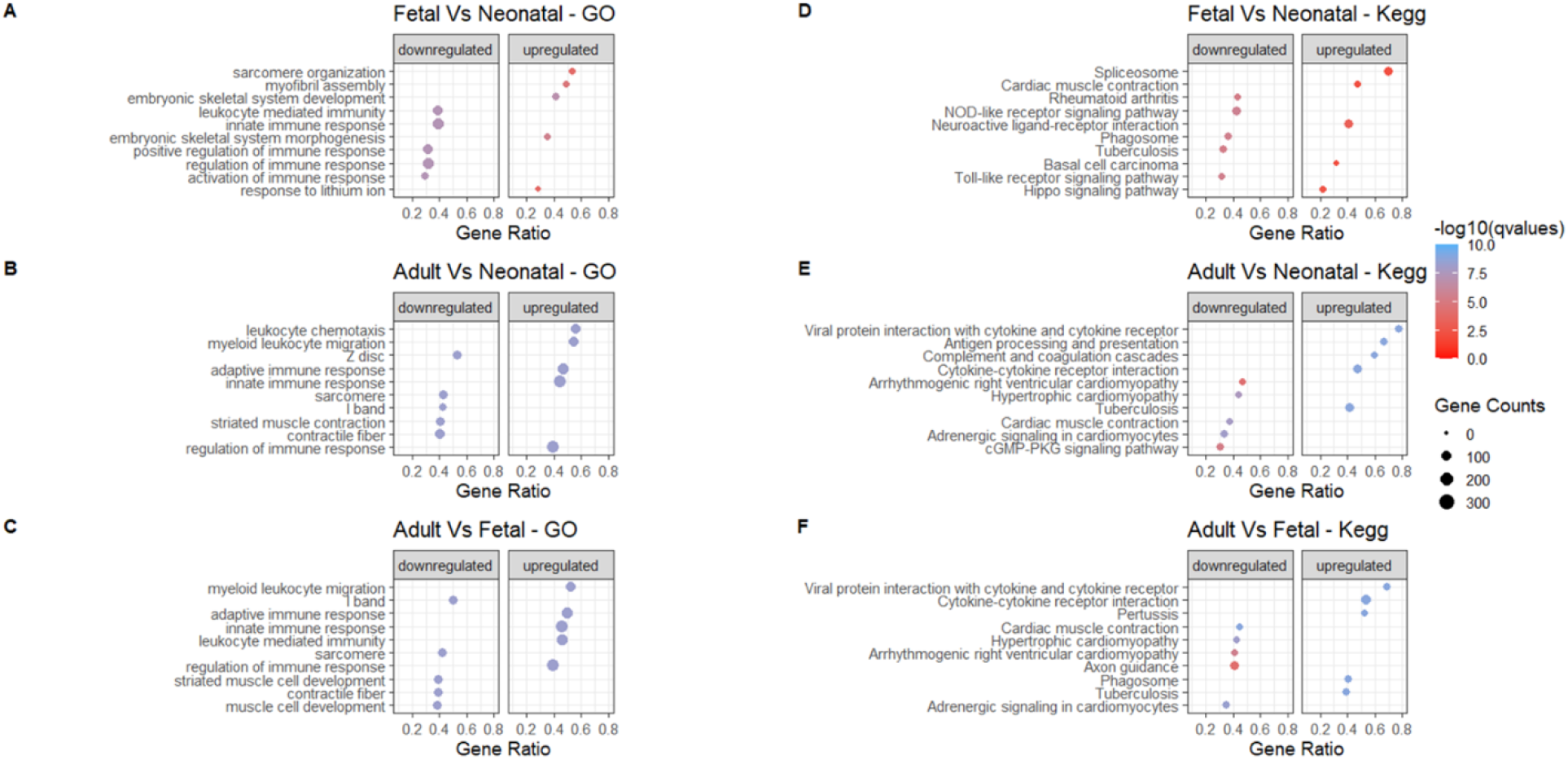
Analysis of functionally enriched GO and KEGG gene sets comparing three ages of cardiac fibroblasts, via Gene Set Enrichment Analysis (GSEA). Dot color is indicative of –log10(q-value), and size is indicative to number of identified core enriched genes. Gene ratio is ratio of identified core enriched genes vs the total number of genes associated with that curated gene set. A, B, C) Dot plots indicating top 5 upregulated and downregulated general GO gene sets for fetal vs neonatal, adult vs neonatal, and adult vs fetal, respectively. GSEA analysis cutoff p-value was 0.05. D, E, F) Dot plots indicating top 5 upregulated and downregulated KEGG gene sets for fetal vs neonatal, adult vs neonatal, and adult vs fetal, respectively. GSEA analysis parameters were set to filter out gene sets with less than 3 genes or greater than 500, or functionally enriched gene sets with p-value > 0.05. Graphs produced in the R programming environment using the clusterProfiler software package.

**Table 2:** Ten most significant (based on q-value) GO and KEGG results from GSEA analysis of Fetal vs. Neonatal differential expression.

| GO |  |  |  |  |  |  |  |  |  |  |  |  |
| --- | --- | --- | --- | --- | --- | --- | --- | --- | --- | --- | --- | --- |
| ONTOLOGY | ID | Description | setSize | enrichmentScore | NES | pvalue | p.adjust | qvalues | rank | count | GeneRatio | (-)LOG10(qvalues) |
| 1 BP | GO:0048706 | embryonic skeletal system development | 98 | 0.66782739 | 2.211347 | 3.09E-10 | 1.41E-07 | 1.22E-07 | 1774 | 41 | 0.41836735 | 6.914235563 |
| 2 BP | GO:0048704 | embryonic skeletal system morphogenesis | 77 | 0.673441241 | 2.163555 | 1.20E-08 | 2.97E-06 | 2.57E-06 | 1173 | 27 | 0.35064935 | 5.589746169 |
| 3 BP | GO:0030239 | myofibril assembly | 63 | 0.685639515 | 2.132596 | 1.46E-07 | 2.24E-05 | 1.94E-05 | 1413 | 31 | 0.49206349 | 4.712815643 |
| 4 BP | GO:0045214 | sarcomere organization | 47 | 0.711616313 | 2.115637 | 1.49E-06 | 0.00014 | 0.000121 | 1413 | 25 | 0.53191489 | 3.915725028 |
| 5 BP | GO:0010226 | response to lithium ion | 38 | 0.737469517 | 2.110208 | 2.41E-06 | 0.000205 | 0.000178 | 705 | 11 | 0.28947368 | 3.749588093 |
| 6 BP | GO:0002443 | leukocyte mediated immunity | 269 | -0.542219601 | -2.25599 | 1.00E-10 | 6.20E-08 | 5.37E-08 | 1818 | 105 | 0.39033457 | 7.269805204 |
| 7 BP | GO:0050776 | regulation of immune response | 457 | -0.516923697 | -2.25641 | 1.00E-10 | 6.20E-08 | 5.37E-08 | 1430 | 144 | 0.31509847 | 7.269805204 |
| 8 BP | GO:0050778 | positive regulation of immune response | 334 | -0.535040447 | -2.28475 | 1.00E-10 | 6.20E-08 | 5.37E-08 | 1430 | 107 | 0.32035928 | 7.269805204 |
| 9 BP | GO:0045087 | innate immune response | 372 | -0.535070216 | -2.2875 | 1.00E-10 | 6.20E-08 | 5.37E-08 | 2068 | 144 | 0.38709677 | 7.269805204 |
| 10 BP | GO:0002253 | activation of immune response | 174 | -0.5822796 | -2.30488 | 1.00E-10 | 6.20E-08 | 5.37E-08 | 1254 | 51 | 0.29310345 | 7.269805204 |
| KEGG |  |  |  |  |  |  |  |  |  |  |  |  |
| ID | Description | setSize | enrichmentScore | NES | pvalue | p.adjust | qvalues | rank | count | GeneRatio | (-)LOG10(qvalues) |  |
| rno04260 | Cardiac muscle contraction | 70 | 0.557570837 | 1.774523 | 0.000675 | 0.008918 | 0.006495 | 2372 | 33 | 0.47142857 | 2.187440803 |  |
| rno05217 | Basal cell carcinoma | 44 | 0.597575616 | 1.74177 | 0.001055 | 0.01126 | 0.0082 | 1309 | 14 | 0.31818182 | 2.086175282 |  |
| rno04390 | Hippo signaling pathway | 128 | 0.488247152 | 1.68712 | 0.000218 | 0.0048 | 0.003496 | 1309 | 28 | 0.21875 | 2.456440873 |  |
| rno04080 | Neuroactive ligand-receptor interaction | 192 | 0.462846994 | 1.672178 | 3.42E-05 | 0.000943 | 0.000687 | 1704 | 78 | 0.40625 | 3.163098363 |  |
| rno03040 | Spliceosome | 113 | 0.486982499 | 1.659018 | 0.000701 | 0.008918 | 0.006495 | 4787 | 79 | 0.69911504 | 2.187440803 |  |
| rno05152 | Tuberculosis | 137 | -0.530667627 | -2.01828 | 1.02E-07 | 6.33E-06 | 4.61E-06 | 1595 | 45 | 0.32846715 | 5.336554399 |  |
| rno04145 | Phagosome | 131 | -0.535473922 | -2.03847 | 4.06E-08 | 4.04E-06 | 2.95E-06 | 1964 | 48 | 0.36641221 | 5.530908671 |  |
| rno04621 | NOD-like receptor signaling pathway | 130 | -0.555866396 | -2.11026 | 7.46E-09 | 2.47E-06 | 1.80E-06 | 2242 | 55 | 0.42307692 | 5.745279883 |  |
| rno05323 | Rheumatoid arthritis | 65 | -0.641598006 | -2.1916 | 1.15E-07 | 6.33E-06 | 4.61E-06 | 1520 | 28 | 0.43076923 | 5.336554399 |  |
| rno04620 | Toll-like receptor signaling pathway | 77 | -0.6261727 | -2.21556 | 4.89E-08 | 4.04E-06 | 2.95E-06 | 1096 | 24 | 0.31168831 | 5.530908671 |  |

Core enriched genes in these datasets included pro-inflammatory chemokines such as *Ccl3, Ccl4, Ccl11,* and immune-associated receptors such as *Ddx58* and *Lbp.* This downregulation in immune-associated ontologies was supported by KEGG results, which indicated downregulation of Nod-like receptor signaling pathways and Toll-like receptor signaling pathways (37) (Fig. 6D, Table 2, Additional File 2).

Upregulated GO terms in FCFs vs NCFs included developmental terms such as “embryonic skeletal system morphogenesis,” and cardiac tissue-associated ontologies including “myofibril assembly” and “sarcomere organization,” (Fig. 6A) supported by KEGG pathways including “cardiac muscle contraction” and “Hippo signaling pathway.” Identified core enriched genes included homeobox proteins (*Hoxb5/7/8, Nkx2.5),* T-box transcription factors (*Tbx1/15*,) and several cardiac tissue proteins including *Actc1, Actn2,* and *Myh6* (Fig 6D, Table 2, Additional File 2).

Similar patterns emerged in analysis of adult CFs compared to both neonatal CFs and fetal CFs. Differential expression in adult vs neonatal and adult vs fetal indicated upregulation of ontology terms associated with adaptive and innate immune response, leukocyte function, and regulation of immune response (Fig 6B & 6C) and in both cases, KEGG pathways including “cytokine-cytokine receptor interaction,” and “viral protein interaction with cytokine and cytokine receptor” were upregulated (Fig 6E & 6F). This was supported by examination of core enriched genes: an array of chemokines including *Ccl3, Ccl4, Ccl7, Ccl11 and Ccl22* were identified as enriched in both adult vs neonatal (Table 3, Additional File 4) and adult vs fetal (Table 4, Additional File 6) analyses, suggesting significant variances in chemokine expression with respect to age.

**Table 3:**
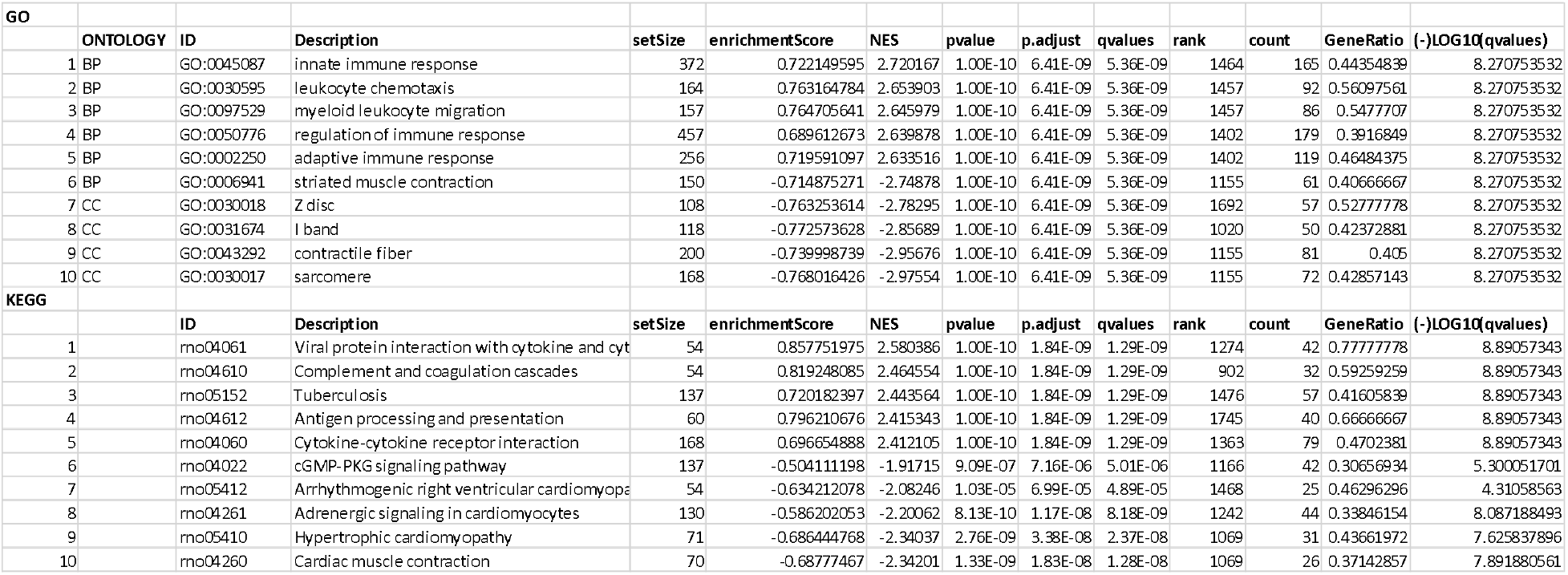
Ten most significant (based on q-value) GO and KEGG results from GSEA analysis of Adult vs. Neonatal differential expression.

**Table 4:**
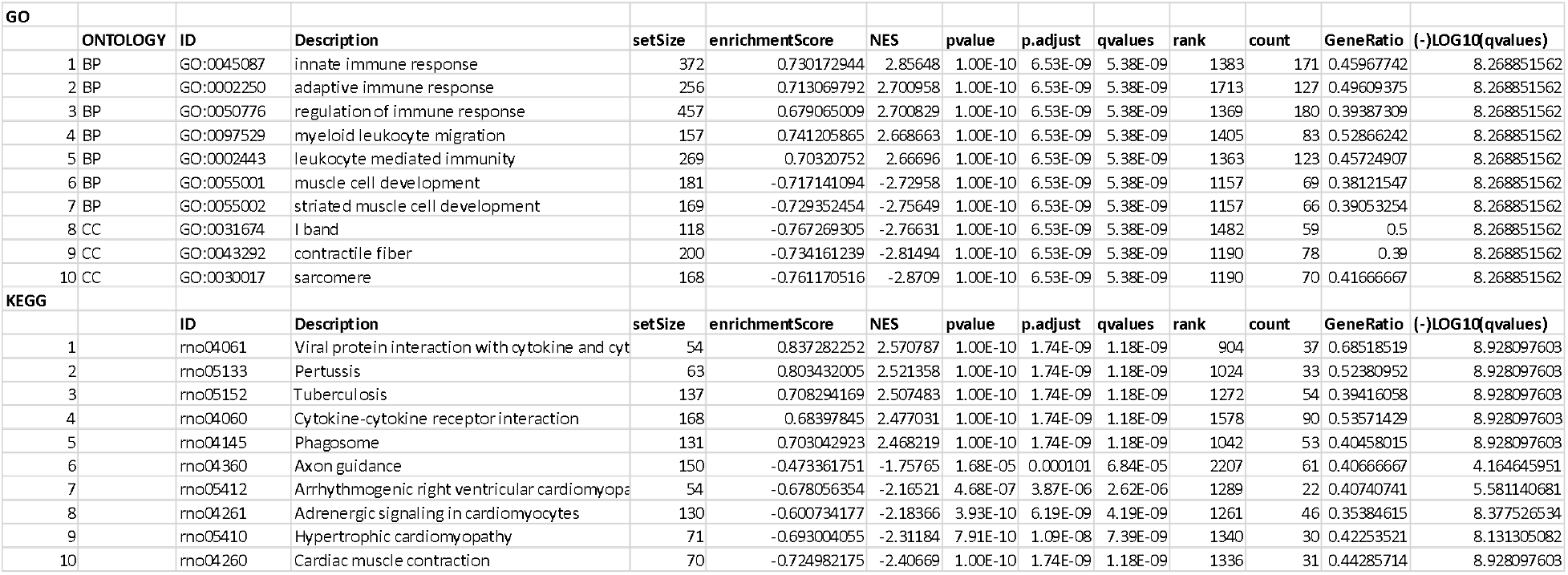
Ten most significant (based on q-value) GO and KEGG results from GSEA analysis of Adult vs. Fetal differential expression.

Downregulation of cardiac function and cardiac development terms were also evident in both comparisons. GO terms indicated as functionally downregulated by the analyses included “contractile fiber,” “striated muscle contraction,” and “sarcomere” in the adult vs neonatal comparison, and similarly “muscle cell development,” and “striated muscle development” were downregulated in the adult vs fetal analysis (Fig 6B & 6C). KEGG terms associated cardiac function including “cardiac muscle contraction” and “adrenergic signaling in cardiomyocytes” were similarly downregulated in both adult vs neonatal and adult vs fetal comparisons. This was supported by examination of core enriched genes for these ontologies, which recapitulated this downregulation of cardiac functional and developmental genes: in both cases genes such as *Myh6/7, Actn1, Actn2, and Actc1,* among several others, were identified (Table 3, Table 4, Additional File 4, Additional File 6).

Taken together, these data underscore a transcriptomic shift from more development-supporting genes to predominantly immune- and inflammatory-specific genes. This shift is evident in comparison of young animal-sourced cardiac fibroblasts (NCF and FCF) to ACFs, but transcriptomic differences also exist between FCFs and NCFs as well.

### RT-qPCR validates differential expression differences in CFs

Reverse transcriptase quantitative PCR (RT-qPCR) was used to analyze FCFs, NCFs, and ACFs in order to validate the RNA-seq results for selected genes. Results show relative expression normalized to GAPDH for ACFs and FCFs, normalized against results from NCFs. The genes evaluated were *Il6, Tbx5, Myh7, Mmp9, Mmp2, and Col1a1. Il6* was selected as Interleukin-6 was shown to be comparatively upregulated in ACF vs NCF (Additional File 5) and is an important inflammatory cytokine. *Tbx5* (T-box transcription factor 5) is a critical cardiac development protein (31) and was observed to be upregulated in FCFs vs NCFs (Additional File 3). *Myh7* (encoding a myosin heavy chain β isoform) was observed to be markedly downregulated in ACFs vs NCFs (Additional File 5). *Mmp2, Mmp9,* and *Col1a1,* encoding matrix metalloproteinase-2 and −9, and collagen type I, alpha-1, are all critical in cardiac matrix regulation and crucial proteins expressed by CFs. Of these, *Mmp9* was shown to be upregulated in ACFs vs. NCFs, while *Mmp2* and *Col1a1* were observed to be marginally downregulated in both ACFs and FCFs compared to NCFs (Fig 7A).

**Figure 7:**
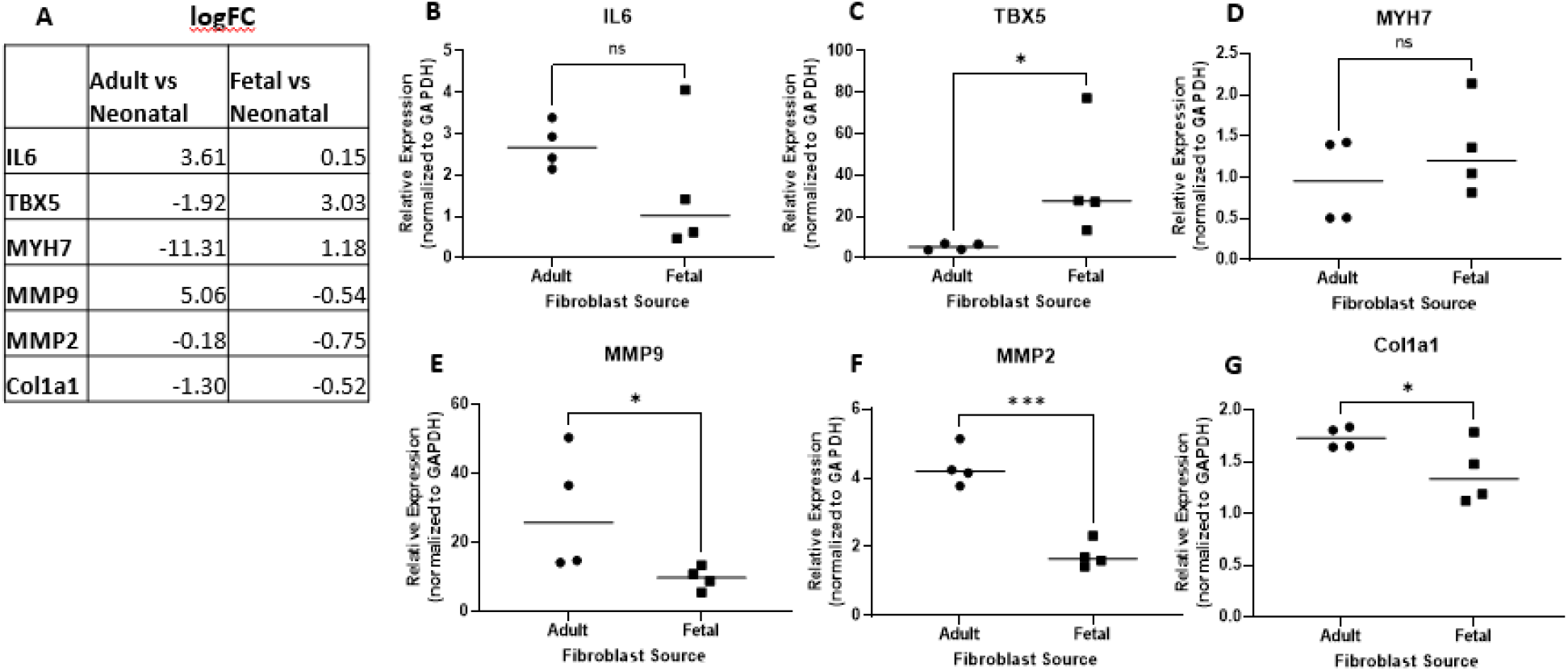
RT-qPCR validation of differentially expressed genes in adult and fetal CFs, with respect to neonatal CFs. A) Log2FC differential gene expression from RNA-seq data for genes selected for validation via qPCR. B) IL6 gene expression in adult CFs vs. fetal CFs, normalized to neonatal CF controls and relative to GAPDH expression. C-G) TBX5, MYH7, MMP9, MMP2, COL1A1 expression levels, respectively, in adult CFs vs. fetal CFs. Statistical analysis: one-tail t-test, with cutoff p-value threshold of 0.05. *=p<0.05, ***=p<0.001, ns = no significance. Error bars indicate standard deviation.

RT-qPCR results recapitulated these observed expressed differences in part: *Il6* had a relative (to NCF, normalized to GAPDH) expression value of 2.72 ± 0.54 (std. dev) in ACFs, vs expression in FCFs of 1.64 ± 1.67 (p=0.1320) (Fig. 5B). *Tbx5* was significantly less expressed (5.33 ± 1.65) in ACFs vs in FCFs (36.38 ± 28.04, p=0.0345) (Fig. 7C) consistent with trends observed in the RNA-seq results, while *Myh7* showed variable, low expression in both cases (ACF: 0.96 ± 0.52, FCF: 1.34 ± 0.58, p=0.1821) (Fig. 5D). *Mmp9* (ACF: 29.06 ± 17.67, FCF: 9.74 ± 3.33, p=0.0376)*, Mmp2* (ACF: 4.33 ± 0.58, FCF: 1.76 ± 0.38, p=0.0002), and *Col1a1* (ACF: 1.73 ± 0.10, FCF: 1.39 ± 0.30, p=0.0387) all had significantly higher expression in ACFs vs FCFs (Fig. 7E, 7F, 7G).

## Discussion

Analysis of the cardiac fibroblast transcriptome in fibroblasts isolated from fetal, neonatal, and adult developmental ages indicates broad differences in gene expression with respect to age. This data offers new insights into the potential functions of CFs at these developmental ages while reinforcing age-specific gene expression differences in CFs and cardiac tissue that have been previously uncovered in the literature. More specifically, this data supports previous conclusions that indicate that fetal CFs present a gene expression profile reflecting functions in support of embryonic growth and cardiac development, whereas their adult CF counterparts upregulate genes more reflective of immune cell communication and trafficking, and cardiac homeostatic maintenance (19, 21, 31, 38, 39). Our data both supports and expands on these studies by incorporating neonatal CFs into the analysis to evaluate fetal-to-neonatal differences. This establishes a clearer timeline of the phenotypic shifts in CFs, from the primarily developmental role of fetal CFs (21, 38), transitioning to a phenotype associated with rapid expansion and maturation in the neonatal heart (40) to an adult phenotype characterized by immune signaling and tissue maintenance (24, 25, 40, 41).

### RNA-seq confirms cardiac fibroblast immune function is age-dependent

Although historically the function of CFs was considered to exclusively ECM maintenance, it is now known these cells possess varied functions fundamental to cardiac function (24, 31, 38), and in particular are vital to cardiac immune and inflammatory processes. In this respect, CFs secrete an array of growth factors, chemokines, and cytokines that facilitate invasion of immune cells such as macrophages and granulocytes (4, 15, 25, 41, 42), including MCP-1, MIP-1α, MIP-1β, IP-10, and RANTES (42, 43).

RNA-seq results in this study reinforces the hypothesis of an immune-regulatory role of cardiac fibroblasts, focusing on how this role varies across organismal development. The lack of a strong immune system in the fetal and neonatal mammalian heart is well-established and hypothesized to contribute to the regenerative wound healing observed in fetal and neonatal mammals (44, 45). It is similarly understood that maturation of the immune system and expression of inflammatory cytokines increases rapidly during post-natal development (46), but an in-depth examination of how the cardiac fibroblast transcriptome changes in an immune context, and how these changes may contribute to immune maturation in the heart has, to our knowledge, not been thoroughly explored. The transcriptomic data presented herein point toward more immune crosstalk and inflammatory signaling in CFs with respect to increasing developmental age, and is worth further investigation and validation. Understanding how CF impact inflammation and immunity at different developmental ages may support the development of regenerative therapies and engineered cardiac tissues by uncovering ways to minimize the tissue inflammatory response and aggressive immune response typical of fibrotic wound repair following injury in adults (11, 47, 48).

We observed an upregulation of genes responsible for macrophage invasion (*Ccl3/Ccl4*) (49) (Table 3, Table 4, Additional File 5, Additional File 7) with respect to developmental age, in addition to increases in the expression of inflammatory mediators (*Il-1β, Il-6)* in adult CFs versus neonatal and fetal CFs (Additional File 5, Additional File 7). Further, gene ontology analysis results support the hypothesis that immune and inflammatory gene expression differences exist between fetal and neonatal CFs, suggesting cardiac immune maturation progresses rapidly post-birth. These included fetal downregulation (compared to neonatal and adult CFs) of immune response-associated gene ontology terms (Fig 6A), downregulation of Nod-like and Toll-like receptor signaling pathways (Fig 6D), which have both been established as expressed in cardiac fibroblasts and are implicated in triggering an immune response to pathogen-associated molecular patterns (15, 42). We also observed a comparative upregulation of chemokines in adult CFs versus fetal CFs, including *Ccl3/4* immune-associated genes like *Ddx58* (an innate immune response receptor essential in identifying viral-infected cells (50) and *Lbp,* which binds to bacterial lipopolysaccharides and initiates an immune response (51) (Table 1).

### Cardiogenic developmental/regenerative potential of cardiac fibroblasts is rapidly lost post-birth

Cardiac fibroblasts, like their counterparts in other bodily tissues, play a vital role in tissue morphology, extracellular matrix regulation and upkeep, mechanical support, and inflammatory response *in vivo* (15, 19, 26, 41, 52, 53). Beyond this, CFs are also implicated in the development and maturation of the heart *in utero* and assist in driving cardiomyocyte proliferation in that regard. This was well-established in work by Ieda *et al.* in 2008 (25), in studies that determined that fibroblast-derived proteins including fibronectin and periostin stimulated cardiomyocyte proliferation via β1-integrin signaling.

It is well-understood that cardiomyocyte proliferation and cardiac regenerative wound healing diminishes rapidly post-birth (22, 24, 54), and is reflected in a reduced regenerative potential in cardiac fibroblasts, which begin to express a more pro-fibrotic phenotype (3, 55, 56). Our study reinforces the age-dependent transition to a fibrotic phenotype in CFs, and suggests that it begins rapidly after birth, as we observed significant downregulation of several immune-associated GO and KEGG pathways in fetal CFs versus neonatal CFs (Fig 6A & 6D, Additional File 2), implying neonatal CFs begin to upregulate genes necessary to mount significant immune recruitment to initiate an inflammatory response immediately after birth. Our data also indicated functional upregulation of genes associated with muscle cell development in fetal and neonatal CFs versus adult CFs (Fig 6B & 6C, Additional File 4, Additional File 6), which may support the conclusion that early-development CFs play a direct role in the functional development of cardiac tissue. This was particularly evident in FCFs, within which development-associated transcription factors including *Tbx4, Hoxb5,* and *Hoxb7* were upregulated, and ontological terms associated with morphogenesis and development were more functionally upregulated than their neonatal counterparts (Fig 6A, 6D, Table 1, Additional File 1). This builds upon previous work investigating transcription factors in cardiac fibroblasts, which concluded that cardiogenic genes including *Tbx20* and *Gata4* expressed by adult fibroblasts contribute to cardiac repair (31, 57).

The presence of cardiomyocyte structural genes including *Actc1, Actn2, Myh6,* and *Myh7* was also observed to be upregulated in FCFs and NCFs versus ACFs, which was surprising as these proteins are conventionally considered cardiomyocyte-specific (57, 58). There is evidence CFs, particularly fetal-derived, express these genes at low levels: a previous microarray assay of fetal versus adult mouse cardiac fibroblast gene expression indicated similar differential expression of these genes with expression levels consistent with what was observed in this study, specifically upregulation of *Actc1, Actn2, Myh6,* and *Myh7* in fetal CFs versus adult CFs (25).

Differences in CF expression of ECM and ECM-associated proteins is also strongly linked to the transition to pro-fibrotic wound healing in the heart. In previous studies of fetal versus adult cardiac fibroblasts, gene expression analysis indicated enrichment of fibronectin (*Fn*), collagen (*Col*) genes, heparin-binding EGF-like growth factor (HBEGF), periostin (*Postn),* and tenascin C (*Tnc*) in fetal cardiac fibroblasts but not adult cardiac fibroblasts, an expression profile associated with increased cardiomyocyte proliferation and regenerative healing (21, 25, 59). Within our differential expression analysis, we observed an upregulation of *Col9a1, Col26a1,* and *Col4a4,* and *Tnc* in fetal CFs vs neonatal CFs (Additional File 3). Of the collagens, both Col9 (60) and Col4 (61) have potential links to cardiac development. *Tnc*, the gene for protein tenascin-C, is predominantly expressed in embryonic tissue and implicated in tissue morphogenesis and remodeling via modulation of cell adhesion to ECM substrates (62). Taken together, this data provides more evidence that the loss a regenerative phenotype in CFs occurs rapidly after birth.

### Study Limitations

This study contains several potential experimental limitations that should be addressed by future investigations. This study exclusively explores the transcriptomic profile of cardiac fibroblasts, and direct evaluations of genes identified must be performed to identify specific factors responsible for the phenotypic differences in CFs hypothesized. The protocols used to isolate cardiac fibroblasts from its resident tissue differed slightly across developmental ages: specifically, while neonatal and adult CFs were from ventricular tissue, fetal CFs were isolated from the whole fetal heart, which was necessary to produce a sufficient cell yield. However, cardiac fibroblast phenotype can vary with respect to location in the heart (63), and incorporation of location-dependent CF RNA-seq analyses would likely be of value. Further, additional single-cell RNA-seq associated studies may be valuable to identify crucial cell-cell heterogeneity associated with CFs, and how this likely differs across developmental age and may be partly responsible for observations presented in this study (24, 32, 64). We also note that the qPCR work was completed with fibroblasts cultured *in vitro* at low passage numbers, versus the RNA-seq analysis which was done with confluent P0 CFs. This could explain why the gene expression in the qPCR study is not entirely consistent with the RNA-seq.

## Conclusion

This transcriptomic analysis of the rat cardiac fibroblast at three discrete developmental ages (fetal, neonatal, and adult) using RNA-seq provides significant insights into CF transcriptional heterogeneity. Immune function-related transcriptomic evaluations presented herein suggest age-specific CF heterogeneity may be partly responsible for differences in cardiac inflammation and immune response observed across developmental ages *in vivo.* The results reinforce established knowledge of fibroblast phenotypic heterogeneity across developmental ages, including marked upregulation of cytokines and chemokines in adult CFs vs fetal and neonatal CFs, and an age-dependent transition to a more pro-fibrotic, pro-inflammatory gene expression profile, versus the pro-regenerative gene expression observed in fetal and early-neonatal CFs. The results further suggest that this transition away from a developmental, pro-regenerative phenotype becomes acute quickly after birth.

## Methods

### Heart harvest from fetal, neonatal, and adult rats

All animal procedures were performed in accordance with the NIH Guide for the Care and Use of Laboratory Animals and approved by the Institutional Animal Care and Use Committee at Tufts University. Rat hearts harvested for cell isolation were from fetal, neonatal, and adult Sprague-Dawley rats. Pregnant dams (~3 months old) and adult rats (~7 weeks old) were deeply anesthetized with 3-5% isoflurane and euthanized via heart removal. Fetal pups (embryonic day 18, E18) were isolated from the uterus, euthanized by conscious decapitation and hearts were isolated. Neonatal pups (postnatal day 1, P1) were euthanized by conscious decapitation prior to heart harvest. Hearts were stored in an ice-cold solution of sterile phosphate-buffered saline (PBS) with 20 mM glucose and immediately used for cell isolation, as described below.

### Primary cardiac fibroblast isolation and culture

Cardiac fibroblasts were isolated from fetal, neonatal, and adult rat hearts following previously described methods (65). After euthanasia and heart harvest, adult heart left ventricles were separated from the heart and minced to pieces less than 1 mm^3^ in size. Fetal and neonatal hearts were minced whole. Minced tissue underwent 3-5 serial digestions in collagenase type 2 (Worthington Biochemical Corp, Lakewood, NJ) in sterile PBS with 20 mM glucose, and passed through a 40 μm cell strainer and centrifuged. CFs were separated from other cardiac cells via collection as fast-adhering cells in a 1-hour pre-plating in Dulbecco’s Modified Eagle Medium (DMEM) with 10% fetal bovine serum (FBS), 1% penicillin-streptomycin (Invitrogen), and 0.1 mM ascorbic acid, followed by removal of nonadherent cells and reapplication of media (3, 25). Fibroblasts for RNA-seq were expanded to confluency in culture and used immediately for RNA isolation. CFs for all other experiments were used at passage 2-4.

### Immunofluorescence

Immunofluorescent imaging was used to validate that the cells used in culture were predominantly cardiac fibroblasts, and not from another nonmyocyte cell lineage. To confirm this, cells were stained for vimentin (a broad marker for fibroblasts but also expressed in endothelial cells (31, 32)), and either CD31 or CD45 to identify endothelial or hematopoietic cells, respectively (30, 31). CFs at passage 2-4 were seeded at a density of 50,000 cells/cm^2^ for 24 hours, and then fixed for 10 minutes with ice-cold 4% paraformaldehyde. Cells were permeabilized with 0.05% Triton X-100 in PBS for 10 minutes at room temperature, rinsed in PBS, and blocked with 5% normal donkey serum and 1% bovine serum albumin (BSA) in PBS overnight at 4 °C. After blocking, cells were incubated with primary antibodies, against vimentin (Cell Signaling Technology, #5741, dilution 1:200), CD31 (Abcam, ab119339, dilution 1:500), and CD45 (Abcam, ab10558, dilution 1:250) diluted in 1% BSA in PBS overnight, again at 4 °C. Cells were rinsed in PBS with 0.1% Tween-20 (PBST) before incubation with fluorescently labeled secondary antibodies (ThermoFisher Donkey Anti-Rabbit 488-A21206, Donkey Anti-Mouse 488-A10037) for two hours, followed by Hoescht (10 μg/mL) and phalloidin (Invitrogen 588-A12380, diluted according to manufacturer’s instructions, ~125ug/mL) for 30 minutes at room temperature. Cells were rinsed in PBST again and immediately imaged on a Keyence BZ-X710 fluorescent microscope.

### RNA Preparation for Sequencing

P0 neonatal, fetal, and adult Sprague Dawley rat cardiac fibroblasts were cultured to confluency in T75 flasks, in serum-containing media with 0.1 mM ascorbic acid. Upon reaching confluent density, the cells were lysed in 1mL of ice-cold TRIzol reagent (Invitrogen) and RNA was isolated via manufacturer’s instructions: briefly, 200uL of chloroform was added to each mL of TRIzol, mixed vigorously, incubated on ice for 2-5 minutes, and centrifuged at 4C at 12,000G for 15 minutes. The aqueous upper layer containing RNA was isolated and purified of chemical contaminants and DNA via processing in a SurePrep RNA Cleanup and Concentration Kit, and treatment with ThermoScientific RapidOut DNA Removal Kit, both following manufacturer’s protocols. RNA was then quantified on a Nanodrop 2000 UV-vis spectrophotometer (ThermoFisher Scientific) and stored at −80C until further use.

### RNA Sequencing

Purified RNA samples from fetal, neonatal, and adult rat hearts (n = 3 for each age group) were delivered to the Tufts University Genomics Core Facility for quality certification, cDNA library construction (Illumina TruSeq Stranded mRNA), and sequencing. Sequencing was performed on the Illumina HiSeq 2500 platform (50-bp single-end lane). Sequencing data in fastq format can be accessed in the Gene Expression Omnibus database (GEO) with the accession number GSE162277. The alignment of sequenced reads to the rat reference genome (Rn6 rat genome from the UCSC Genome Browser) was performed in several steps. First, sequenced data was adapter-trimmed using Trimmomatic 0.39 (66). Pre- and post-trimming, raw sequence data was quality validated using FASTQC (67). Trimmed and quality-assessed data was loaded into STAR (v. 2.5.2b) (68) for read alignment to the Rn6 rat genome. STAR-aligned reads stored as BAM files loaded into featureCounts (69) for read counting. Raw counts were imported into R for differential gene expression analysis.

### Differential Gene Expression Analysis

Raw counts were imported into R for gene expression analysis, using the *DESeq2* package, using the workflow described in the DESeq2 manual (70). Briefly, DESeq2 normalizes counts via a median of ratios method, in which counts are divided by sample-specific size factors determined by the median ratio of gene counts, relative to the geometric mean per gene (71). Principal component analysis (PCA) was performed with the plotPCA function in R, using natural log-transformed expression data from DESeq2. Normalized gene expression data from the three groups was output into a hierarchical clustering heatmap with the *pheatmap* package in R. These analyses were done to check for batch effects, and to broadly evaluate the gene expression data across the three age groups. Calculation of number of reads per sample (in millions of reads), correlation between samples, analysis of loadings of individual genes in the principal components, and clustering of individual genes with respect to principal components was performed in pcaExplorer (72), which utilizes the DESeq2-transformed data in its gene set analyses.

The neonatal cardiac fibroblast group (NCF) was assigned as the control condition for analysis, and differential expression comparisons to adult cardiac fibroblasts (ACF) and fetal cardiac fibroblasts (FCF) were calculated, although the comparison case ACF vs FCF was also analyzed and included in results. The full datasets pre-ranked by log_2_FC were imported into the *clusterProfiler* package in R (34), which includes methods incorporating Gene Set Enrichment Analysis (GSEA) (33) and analyzed against Gene Ontology (GO) (35) and Kyoto Encyclopedia of Genes and Genomes (KEGG) (36) databases to identify enriched gene sets and biological pathways. GSEA was run with gene set size thresholds of a minimum of 3 and maximum of 500, p-value cutoff of 0.05, and p-value adjustment using the Benjamini-Hochberg method (73). Separately, expression results were filtered against a cutoff log_2_FC and adjusted p-value (abs(logFC) ≥ 1.5, padj < 0.05) to create datasets of significantly differentially expressed genes for downstream evaluation.

### RT-qPCR Gene Expression Validation

CFs at passage 2-4 were lysed and RNA isolated and purified using a SurePrep DNA/RNA/Protein Concentration Kit, following manufacturer’s protocols. RNA was quantified on a Nanodrop and stored at −80C until use. Following RNA purification, RNA was converted to cDNA using a Thermofisher Scientific High Capacity cDNA Reverse Transcriptase Kit and cDNA was stored at −80C until further analysis. Reverse transcriptase quantitative PCR (RT-qPCR) was performed using an Applied Biosystems TaqMan Gene Expression Master Mix, carried out on a BioRad CFX96 Real-Time System using TaqMan PCR primers (ThermoFisher Scientific) according to manufacturer instructions using 50ug of cDNA per reaction. The genes assayed were COL1A1 (Collagen 1 α1 chain, Rn01463848_m1), MMP2 (matrix metalloproteinase 2, Rn01538170_m1), MMP9 (Rn00579162_m1), IL6 (interleukin 6, Rn01410330_m1), myosin heavy chain beta (MYC7, Rn01488777_g1), and GAPDH (Rn01775763_g1). Genes of interest were normalized to GAPDH expression, with comparative C_T_ used for analysis using neonatal cardiac fibroblasts as a control group.

### Statistical Analysis

Statistical analysis of differential expression data was performed entirely in the R programming environment. As stated above, when identifying significantly expressed genes, a cutoff adjusted p-value of 0.05 was applied. For qPCR evaluation, statistical significance was determined via a one-tailed t-test, with significance set at p < 0.05. Statistical analysis and data visualization of qPCR data was done in the GraphPad Prism environment.

## Additional Files

**Additional File 1:** Top 10 most upregulated and most downregulated (log_2_FC) genes in each differential expression comparison, with associated functional descriptions.

**Additional File 2:** Full Lists of GSEA-identified GO and KEGG functionally enriched terms for Fetal vs. Neonatal comparison

**Additional File 3:** Full List of DESeq2-calculated differentially expressed genes in Fetal vs. Neonatal comparison.

**Additional File 4:** Full Lists of GSEA-identified GO and KEGG functionally enriched terms for Adult vs. Neonatal comparison

**Additional File 5:** Full List of DESeq2-calculated differentially expressed genes in Adult vs. Neonatal comparison.

**Additional File 6:** Full Lists of GSEA-identified GO and KEGG functionally enriched terms for Adult vs. Fetal comparison

**Additional File 7:** Full List of DESeq2-calculated differentially expressed genes in Adult vs. Fetal comparison.

## Acknowledgements

This work was supported by the United States Department of Defense (W81XWH-16-1-0304 to LDBIII), the National Science Foundation (NSF#1603524 to LDBIII) the American Heart Association (18PRE33960362 to LRP), the NIH (R00-CA207866-05 to MJO), and Tufts University (Start-up funds from the School of Engineering to MJ.O and a Breast Cancer Alliance Young Investigator Grant to MJO and TL)

## Abbreviations

cDNA: complementary deoxyribonucleic acid
CF: cardiac fibroblast
CM: cardiomyocyte
DAPI: 4’,6-diamidino-2-phenylindole
DDR2: discoidin domain receptor 2
DNA: deoxyribonucleic acid
ECM: extracellular matrix
FBS: fetal bovine serum
GAPDH: glyceraldehyde 3-phosphate dehydrogenase
GO: gene ontology
IL: interleukin
KEGG: Kyoto Encyclopedia of Genes and Genomes
logFC: log fold change
LPS: lipopolysaccharide
MMP: matrix metalloproteinase
MYH: myosin heavy chain
mRNA: messenger ribonucleic acid
PBS: phosphate buffered saline
PCA: principal component analysis
RNA: ribonucleic acid
RNA-seq: RNA sequencing
RT-qPCR: reverse transcriptase quantitative polymerase chain reaction

