## Additional File 1 for "RNA-sequencing indicates immune cell signaling and inflammatory gene expression in cardiac fibroblasts increases with developmental age"

Top Ten Most Upregulated Genes in All Three Differential Expression Comparisons

| **Fetal Vs Neo Genes** | **log2FoldChange** | **Description** | **Ref** | **Adult vs Neo Genes** | | **log2FoldChange** | **Description** | **Ref** | **Adult vs Fetal Genes** | **log2FoldChange** | **Description** | **Ref** |
| --- | --- | --- | --- | --- | --- | --- | --- | --- | --- | --- | --- | --- |
| Upk3bl | 5.91919 | Uroplakin 3b-like protein, implicated in epithelial development | (1) | Ccl22 | | 10.2823 | C-C motif chemokine 22, involved in immune cell signaling in inflammation | (2) | Clec10a | 11.5862 | C-type lectin receptor family member 10A, an immune-associated endocytic receptor | (3) |
| Tbx4 | 5.71153 | T-box 4, transcription factor in embryonic development | (4, 5) | Cd74 | | 9.17608 | Cell surface receptor for inflammatory cytokine MIF | (6) | Serpina3n | 11.0774 | Serine protease inhibitor with roles roles including wound healing, complement cascade, ECM remodeling | (7) |
| Hoxb7 | 5.66558 | Homeobox 7, cardiac development-associated transcription factor | (8) | Ccl17 | | 8.78364 | C-C motif chemokine 17, ligand for CCR4, involved in immune cell trafficking | (9) | Asgr2 | 10.4007 | Encodes subunit of asialoglycoprotein receptor. Mediates endocytosis of glycoproteins. | (10) |
| Igfbp5 | 5.59857 | Insulin-like growth factor-binding protein 5, implicated in regulation of tissue growth/development | (11) | Asgr2 | | 8.23906 | Encodes subunit of asialoglycoprotein receptor. Mediates endocytosis of glycoproteins. | (10) | Ccl22 | 10.3674 | C-C motif chemokine 22, involved in immune cell signaling in inflammation | (2) |
| Myog | 5.50549 | Myogenin, transcriptional activator necessary in myogenesis | (12) | Serpina3n | | 8.08036 | Serine protease inhibitor with roles roles including wound healing, complement cascade, ECM remodeling | (7) | Ccl17 | 9.92208 | C-C motif chemokine 17, ligand for CCR4, involved in immune cell trafficking | (9) |
| Dhrs9 | 5.36297 | Dehydrogenase/  reductase 9 – metabolic enzyme | (13) | Mmp7 | | 7.74427 | Matrix metalloproteinase-7, associated with LV remodeling post-MI | (14, 15) | Pla2g2a | 9.60937 | Secretory phospholipase A2 group IIA, associated with progression of CHD, inflammation, and thrombosis | (16, 17) |
| Tenm4 | 5.19355 | Teneurin transmembrane protein 4, implicated in mesoderm induction | (18) | Pla2g2a | 7.51227 | | Secretory phospholipase A2 group IIA, associated with progression of CHD, inflammation, and thrombosis | (16, 17) | Akr1c14 | 9.35403 | 3α-hydroxysteroid dehydrogenase, enzyme which catalyzes formation of various neurosteroids | (19) |
| Gldn | 5.1302 | Gliomedin, necessary in peripheral nervous system development | (20) | RT1-Ba | | 7.50239 | Rano class II histocompatibility antigen, B alpha chain | (21) | Ccl4 | 9.26584 | MIP-1β, macrophage inflammatory protein 1-beta, macrophage chemoattractant | (22) |
| Hoxb5 | 5.10057 | Homeobox 5, vascular development-associated transcription factor | (23) | Irg1 | | 7.48663 | Immune responsive gene 1, mitochondrial enzyme that produces itaconate in inflammatory conditions, linked to inflammatory regulation | (24, 25) | Ccl3 | 9.02789 | MIP-1α , macrophage inflammatory protein 1-alpha, macrophage chemoattractant | (22) |
| Mfap2 | 5.05297 | Microfibrillar-associated protein 2, ECM protein | (26) | Chi3l1 | | 7.47072 | Chitinase 3-like 1, inflammatory promoter implicated in macrophage activation, tumor growth | (27) | LOC24906 | 9.02639 | RoBo-1, implicated in bone & extracellular matrix remodeling | (28, 29) |

Top Ten Most Downregulated Genes in All Three Differential Expression Comparisons

| **Fetal Vs Neo Genes** | **log2FoldChange** | **Description** | **Ref** | **Adult vs Neo Genes** | **log2FoldChange** | **Description** | **Ref** | **Adult vs Fetal Genes** | **log2FoldChange** | **Description** | **Ref** |
| --- | --- | --- | --- | --- | --- | --- | --- | --- | --- | --- | --- |
| Akr1c14 | -5.5665 | 3α-hydroxysteroid dehydrogenase, enzyme which catalyzes formation of various neurosteroids | (19) | Myh7 | -11.308 | β-myosin heavy chain, myosin isoform in muscle fibers | (30) | Myh7 | -12.484 | β-myosin heavy chain, myosin isoform in muscle fibers | (30) |
| LOC24906 | -4.8024 | RoBo-1, implicated in bone & extracellular matrix remodeling | (28, 29) | Actc1 | -11.197 | Cardiac α-actin 1, actin isoform predominantly expressed in cardiac tissue | (31) | Actc1 | -12.317 | Cardiac α-actin 1, actin isoform predominantly expressed in cardiac tissue | (31) |
| Figf | -4.5869 | c-fos-induced growth factor, a vascular and endothelial growth factor | (32) | Myh6 | -10.978 | α-myosin heavy chain, myosin isoform | (33) | Myh6 | -11.807 | α-myosin heavy chain, myosin isoform | (33) |
| Mrgprf | -4.4032 | Mas-related G protein-coupled receptor F | (34, 35) | Nppa | -10.745 | Encodes atrial natriuretic peptide, cardiac hormone involved in cardiac development | (36) | Myl2 | -11.492 | Myosin light chain-2, important myosin isoform implicated in contractile function during embryonic development | (37) |
| Clec10a | -4.1869 | C-type lectin domain family 10, member A – endocytic receptor on antigen-presenting cells | (3) | Myl2 | -9.9243 | Myosin light chain-2, important myosin isoform implicated in contractile function during embryonic development | (37) | Nppa | -11.403 | Encodes atrial natriuretic peptide, cardiac hormone involved in cardiac development | (36) |
| Cpxm2 | -4.011 | Carboxypeptidase X, M14 family member 2 – development-associated zinc-dependent peptidase | (38) | Myl3 | -9.3086 | Myosin light chain-3, a ventricular myosin isoform | (39) | Tnni1 | -10.609 | Slow skeletal muscle TnI (troponin 1), TnI isoform present in embryonic hearts | (40) |
| Slc47a1 | -3.9013 | MATE1, multidrug and toxic compound extrusion transporter-1 | (41) | Sh3bgr | -8.9804 | SH3 domain binding glutamate-rich protein implicated in cardiac development | (42) | Sh3bgr | -10.414 | SH3 domain binding glutamate-rich protein implicated in cardiac development | (42) |
| Dpysl4 | -3.8462 | Dihydropyrimidinase-related protein 4, implicated in neuron development, differentiation of epithelial cells | (43) | Tnni1 | -8.979 | Slow skeletal muscle TnI (troponin 1), TnI isoform present in embryonic hearts | (40) | Myl3 | -10.376 | Myosin light chain-3, a ventricular myosin isoform | (39) |
| Mnda | -3.77 | Myeloid nuclear differentiation antigen | (44) | Csrp3 | -8.7669 | Cysteine and glycine rich protein 3, implicated in actin-remodeling and maintenance of cytoskeleton | (45) | Csrp3 | -9.9093 | Cysteine and glycine rich protein 3, implicated in actin-remodeling and maintenance of cytoskeleton | (45) |
| Scn7a | -3.6793 | Sodium channel protein type 7 subunit alpha, encodes sodium channel activated by high extracellular sodium concentration about 150 mM | (46) | Acan | -8.4899 | Aggrecan, proteoglycan and ECM protein present during cardiac development | (47) | H19 | -9.6351 | Long-coding RNA (lncRNA) upregulated in fetal cardiac tissue, downregulated after birth | (48) |
